# Generation of Vascularized Brain Organoids to Study Neurovascular Interactions

**DOI:** 10.1101/2022.01.04.474960

**Authors:** Xin-Yao Sun, Xiang-Chun Ju, Yang Li, Peng-Ming Zeng, Jian Wu, Li-Bing Shen, Yue-Jun Chen, Zhen-Ge Luo

## Abstract

The recently developed brain organoids have been used to recapitulate the processes of brain development and related diseases. However, the lack of vasculatures, which regulate neurogenesis, brain disorders, and aging process, limits the utility of brain organoids. In this study, we induced vessel and brain organoids respectively, and then fused two types of organoids together to obtain vascularized brain organoids. The fused brain organoids were engrafted with robust vascular network-like structures, and exhibited increased number of neural progenitors, in line with the possibility that vessels regulate neural development. Fusion organoids also contained functional blood-brain-barrier (BBB)-like structures, as well as microglial cells, a specific population of immune cells in the brain. The incorporated microglia responded actively to immune stimuli to the fused brain organoids. Thus, the fusion organoids established in this study allow modeling interactions between the neuronal and non-neuronal components *in vitro*, in particular the vasculature and microglia niche.

## Introduction

Recently, brain organoids derived from human pluripotent stem cells (PSCs), including induced PSCs (iPSCs) and embryonic stem cells (ESCs), have been developed to model developmental programs of human fetal brain, recapitulate developmental, psychiatric, and degenerative brain diseases (Amin and Pasca, 2018; Di Lullo and Kriegstein, 2017; Kelava and Lancaster, 2016; Lancaster and Knoblich, 2014). However, the lack of neurovascular system, which is not only required for oxygen and nutrient supply, but also regulates neurogenesis and brain functions (Delgado et al., 2014; Tata et al., 2016; Zhao et al., 2015; Zlokovic, 2011), limits the applications of brain organoids. Thus, vascularization of brain organoids represents one of the most demanded improvements in the field (Di Lullo and Kriegstein, 2017; Giandomenico and Lancaster, 2017; Kelava and Lancaster, 2016; Mansour et al., 2021).

Blood vessels of the vertebrate brain are formed via sequential vasculogenesis and angiogenesis processes, which involve the initial invasion of endothelial cells (ECs) into the neuroepithelium regions via the perivascular plexus, their subsequent coalescence into primitive blood vessels, and growth and remodeling to form a mature vascular network (Lee et al., 2009). ECs are derived from the mesoderm-derived angioblasts (Zadeh and Guha, 2003). It has been shown that ECs can be derived *in vitro* from human PSCs, and these ECs can be potentially useful in engineering artificial functional blood vessels (Harding et al., 2017). However, the generation of complex vascularized organs from PSCs is still challenging, because it depends on the exquisite orchestration of cues from multiple germ layers and the gene expression profiles of ECs are controlled by finely patterned micro-environmental cues during organogenesis (Cleaver and Melton, 2003). Recently, Wimmer et.al reported the generation of self-organizing human blood vessel organoids induced from PSCs, and the application in the study of diabetic vasculopathy (Wimmer et al., 2019). Given the mesodermal origin of ECs and ectodermic origin of neural fates (Nostro et al., 2008; Stern, 2005), one barrier for the generation of vascularized brain organoids is the difficulty in simultaneous application of induction factors for distinct germ layers and cell fates due to mutual repression.

Several strategies have been used to establish the vascularized brain organoids. One strategy took advantage of natural angiogenesis of host blood vessels that sprout and grow into the grafted cerebral organoids, which later exhibited lower cell death rate and enhanced maturation (Mansour et al., 2018). Another approach utilized transcription factor-mediated differentiation of a subset of PSCs into EC-like cells during cerebral organoid induction, whose maturation process was also enhanced (Cakir et al., 2019). Two additional studies have tried co-culture with ECs or their progenitors during cerebral organoid formation (Lopez-Ramirez et al., 2019; Shi et al., 2020). Although grafted brain organoids appeared to have established functional blood vessels (Mansour et al., 2018), none of these methods can form an entirely and integrated vascular network in the cerebral organoids *in vitro*. And they all lacked the functional microglial cells, the only lifelong resident immune cells, which are derived from mesodermal origin (Muffat et al., 2016). In addition, the blood-brain-barrier (BBB), the structure mainly composed of ECs, astrocytes and pericytes, which protects the brain from circulation, is also lacking in the current ectodermal brain organoid models. By co-culture of primary ECs, pericytes and astrocytes, the BBB spheroids were created as an *in vitro* screening platform for brain-penetrating agents (Bergmann et al., 2018; Cho et al., 2017).

Here, we develop an induction approach for brain-specific vascular organoids, which were cultured in medium containing neurotrophic factors at the maturation stage, to obtain cerebrovascular characteristics of the vessel organoids. Interestingly, a large number of microglial cells were induced by this approach along with other types of vascular cells. The vessel organoids were then fused with the cerebral organoids in the Matrigel, leading to the formation of vascularized brain organoids with invasion of microglia, which could be activated upon immune stimuli. Thus, this study invents an advanced strategy that incorporate vascular and microglia into brain organoids, providing a platform for the study of interactions between neuronal and non-neuronal components during brain development and functioning.

## Results

### Generation of the Vessel Organoids (VOrs)

It has been shown that the canonical Wnt signalling is required for development of ESC-derived mesoderm (Lindsley et al., 2006) and the activation of Wnt signalling induces the mesoderm differentiation from human PSCs (Nostro et al., 2008). Considering that EC-generating vascular progenitors (VPs) are derived from mesoderm during embryogenesis (Gupta et al., 2006), we performed guided mesodermal induction of H9 human embryonic stem cells (hESCs), followed by endothelial differentiation. First, we treated 2-day old (D2) embryonic bodies (EBs) from hESCs, which stably expressed GFP, with GSK3 inhibitor CHIR99021 to activate the canonical Wnt signalling for mesoderm induction (Figure 1A).

**Figure 1.**
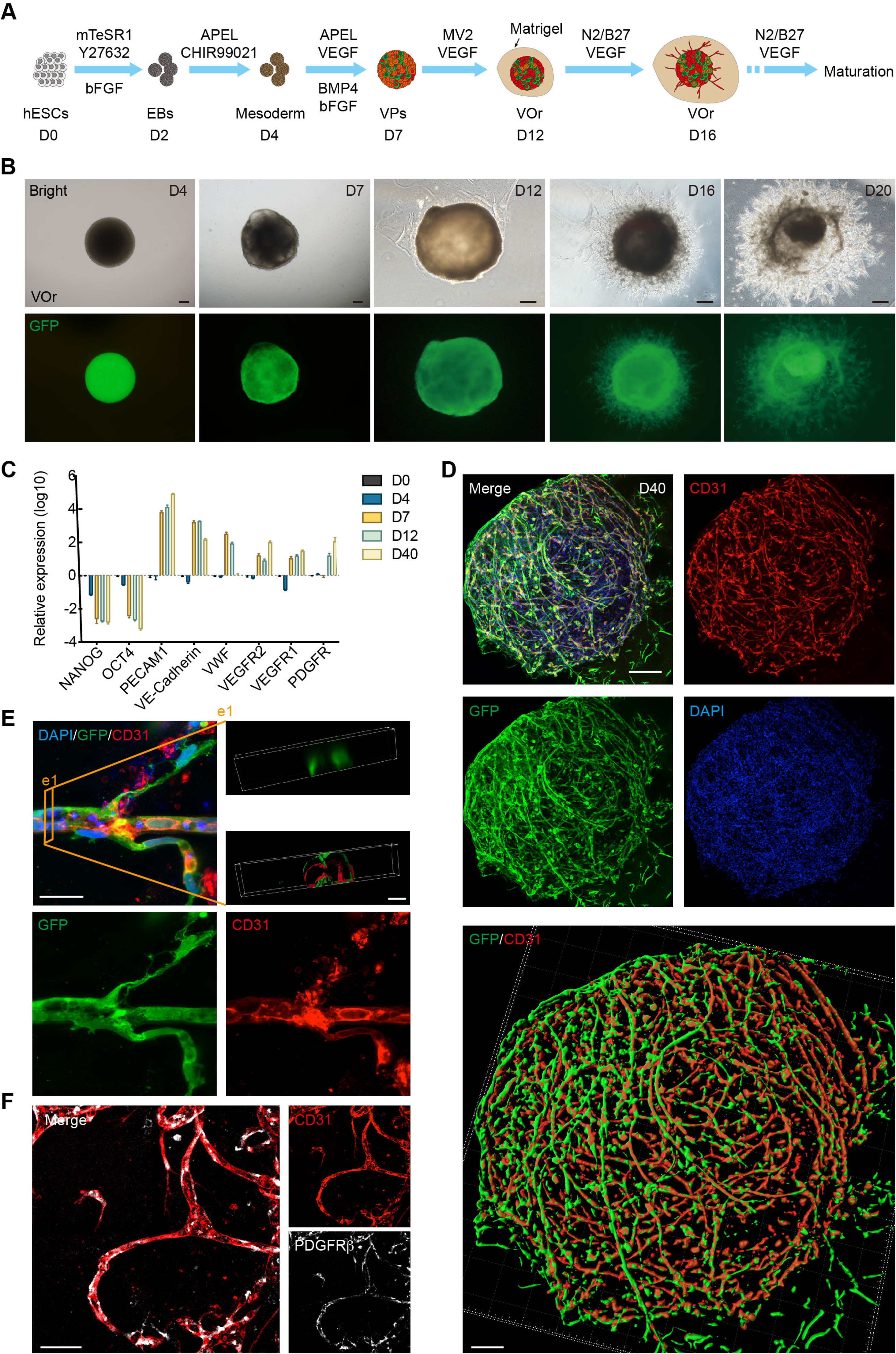
Generation of an *in vitro* model of vessel organoids. (**A**) Schematic view of the methods for generating VOrs from GFP-hESC. EBs: embryonic bodies; VPs: vascular progenitors; VOr: vessel organoid. (**B**) Different developmental stages of VOrs from D4 to D20. Top, right field; bottom, GFP. Scale bar, 200 μm. (**C**) qPCR analysis for expression of stem markers (*NANOG, OCT4*) and vessel markers (*PECAM1, VE-Cadherin, VWF, VEGFR1, VEGFR2, PDGFR*) in developing VOrs, using GAPDH as internal control. Data are presented as mean ± SEM (n = 3 independent experiments). (**D**) Immunostaining of GFP and CD31 in D40 VOrs. Scale bar, 200 μm. Bottom: Imaris reconstruction of VOrs showing integrated vasculature structures. (**E**) Immunostaining of GFP and CD31 for the vascular structures in VOrs. Scale bar, 20 μm. Top right: Section view in VOr showing the lumen structure. (**F**) Immunostaining of CD31 and PDGFRβ for endothelial cells and pericytes, respectively. Scale bar, 50 μm.

After 2 days, the EBs were treated with basic fibroblast growth factor (bFGF), vascular endothelial growth factor (VEGF), and bone morphogenetic protein 4 (BMP4), all of which have been shown to be able to promote VP differentiation into ECs (Cai et al., 2012; Jih et al., 2001). After 3 days, the differentiated ECs were incubated with endothelial medium ECGM-MV2 (shorted as MV2 hereafter) containing VEGF for 5 days for further maturation, and then embedded in Matrigel droplets. At late maturation stages, neurotropic reagents N2 and B27 were added into the maturation medium, which presumably might be able to induce some cell types with specific brain-vessel features (Figure 1A). Notably, vessel-like structures sprouted out from the spheroids (Figure 1B), reminiscent of initial vasculogenesis and angiogenesis. Remarkably, the vessel organoids (VOrs) showed gradual increase in the size during the maturation stage after day 16 (D16), with apparent tubular network characteristics (Figure 1-figure supplement 1A and B).

To verify the cell fates in developing VOrs, we performed quantitative PCR to determine the expression of stemness or vascular-specific genes at different time upon organoid differentiation. As shown in Figure 1C, the stemness markers (*NANOG, OCT4*) showed marked decrease 2 days upon mesoderm induction (D4), whereas the vessel markers (*PECAM1, VE-cadherin, VWF, VEGFR1, VEGFR2 and PDGFRβ*) markedly increased after VP differentiation (D7 and thereafter). In line with this, flow cytometry results revealed the appearance of GFP^+^CD31^+^ ECs after D7, indicating the induction of the endothelial cells (Figure 1-figure supplement 1C).

Morphologically, CD31^+^ ECs in VOrs at D40 showed integrated and complex structures (Figure 1D), and exhibited remarkable vascular branches and tips undergoing angiogenesis-like processes (Figure 1E and Figure 1-figure supplement 1D). The 3D reconstructed cross-section revealed tubular structures in VOrs, reminiscent of vessel lumen (Figure 1E, see top right e1). To determine the connectivity and integrity of vessel-like structures, we injected fluid into the lumens and observed that the hydraulic pressure caused liquid flow and vessel wall expansion without leakage (see Movie Supplement 1).

Notably, PDGFRβ-labeled pericytes that are believed to regulate EC maturation, stabilize vessel wall, and control angiogenesis, were also observed in close contact with ECs undergoing vessel differentiation (Figure 1F). For more details about the vasculature morphology, we used the Angiotool software which had been used as a tool for quantitative vessel analysis (Zudaire et al., 2011). The average vessel length was around 400 mm, the vessel lacunarity was 0.15, and the total number of junctions was about 700-800 per VOr (Figure 1-figure supplement 1E).

To assess the function of ECs in VOrs, we determined the ability to incorporate DiI-acetylated low density lipoprotein (DiI-Ac-LDL), as shown in a previous study (Lehle et al., 2016). We found that VOrs after D14 already had the ability of up-taking DiI-Ac-LDL, whereas ESCs could not (Figure 1-figure supplement 1F). Thus, we have successfully established a fully structured and functional vessel organoid model.

### Cell composition of brain-specific VOrs resembles brain vessels *in vivo*

The vessel system of the brain contains a variety of vascular cell types (Vanlandewijck et al., 2018). To investigate the fidelity of VOrs in recapitulating the cerebrovascular cell types, we performed single-cell RNA sequencing (scRNA-seq) of VOrs at D40 using 10xGenomic chromium system (Macosko et al., 2015; Zilionis et al., 2017). After the quality control data filtering, we analyzed transcriptome of about 7000 single cells, with 200-7000 genes detected per cell and the mitochondrial gene ratio under 5%. The mean reads per cell of two batches of independent samples were highly correlated (Figure 2-figure supplement 1A), indicating negligible batch variance. According to cell type markers of the mice brain vessels identified by single-cell sequencing (He et al., 2018; Vanlandewijck et al., 2018), the cells in VOrs were clustered into 9 main cell types (Figure 2A), including fibroblast (FB), pericyte (PC), proliferative vascular progenitor (MKI67^+^ VP), endothelial cell (EC), smooth muscle cell (SMC), microglia (MG), immune cell (IM) and unknown cluster (Figure 2B). FB accounted for the highest proportion of total cells and MG the lowest (Figure 2-figure supplement 1B). We chose the top 5 highly expressed marker genes of each cluster (Figure 2-figure supplement 1C), and analyzed their expression patterns in each cell type (Figure 2C). Immunostaining showed that the vasculatures in VOrs exhibited positive signals of the SMC marker αSMA, the pericyte marker PDGFRβ, and the EC marker CD31 (Figure 2D and E), confirming the results obtained using scRNA-seq. Meanwhile, the presence of MG-like cells was verified by the staining with specific markers TREM2 and TMEM119 (Figure 2F). Interestingly, DLL4 and EPHB4, which mark the venous and arteries endothelial cells, respectively (Vanlandewijck et al., 2018; Zhao et al., 2018), were found to express only in separate EC populations (Figure 2G and H). This result indicates that ECs in VOrs already underwent spontaneous functional maturation. Immunostaining also confirmed the presence of the venous and arteries EC sub-types (Figure 2-figure supplement 1D and E). Thus, the formed VOrs contained the repertoire of brain-vessel cell types resembling that *in vivo*.

**Figure 2.**
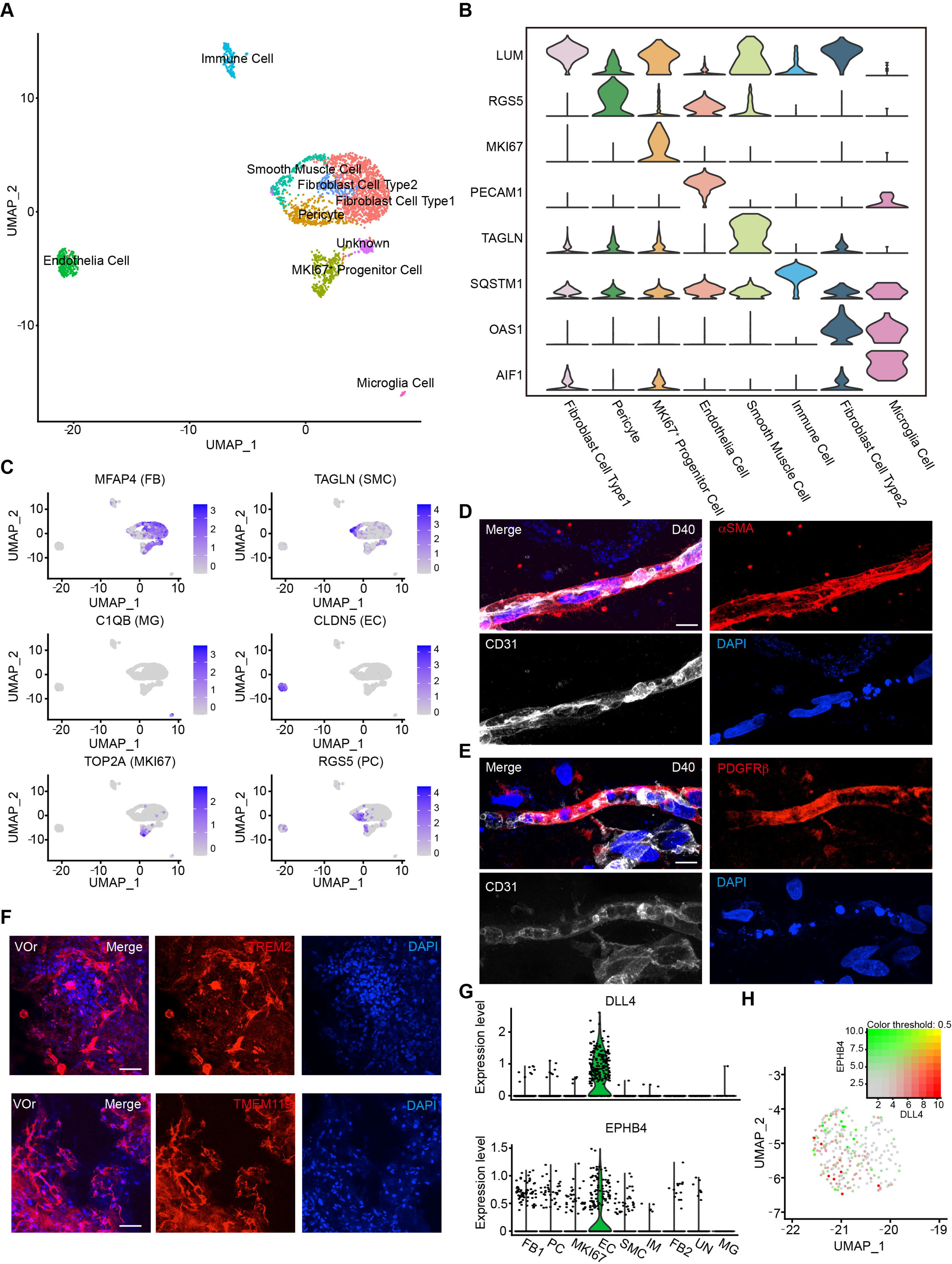
Single-cell transcriptomic analysis of vessel organoids. (**A**) UMAP plot showing the nine major cell types isolated from D40 VOrs. (**B**) Violin plots showing the expression value of the typical markers in each cluster. (**C**) Expression pattern of cell-type specific markers in VOrs. Relative expression level is plotted from gray (low) to blue (high) colors. (**D**) Immunostaining of αSMA for representing the smooth muscle cells in VOrs. Scale bar, 10 μm. (**E**) Immunostaining of PDGFRβ for representing the pericytes in VOrs. Scale bar, 10 μm. (**F**) Immunostaining of microglia markers TREM2 and TMEM119 in VOrs at D40. Scale bar, 50 μm. (**G**) Violin plots showing the expression value of the venous marker EPHB4 and arterial marker DLL4 in EC clusters. (**H**) Expression pattern of arterial and venous markers in EC clusters. Relative expression level is plotted from gray to green (EPHB4) or red (DLL4) colors.

To depict the developmental process of VOrs, we reconstructed the time-course of vascular cell developmental trajectory in Pseudotime (Figure 3A and Figure 3-figure supplement 1A). Five developmental stages and two time points were showed in the trajectory, with stages 1 and 2 representing initial states, stage 3 representing the intermediate state, and stages 4 and 5 the latest (Figure 3-figure supplement 1A and B). Then, we used a panel of markers to annotate the main cell types and found that FB and PC were among the early developed cell types while the EC and MG were among the later ones (Figure 3B-D and Figure 3-figure supplement 1C). It is known that PC and SMC constitute mural cells of blood vessels and it has been difficult to distinguish them because they have similar gene expression profiles (Smyth et al., 2018). Using the developmental trajectory analysis, we found that PC appeared earlier than SMC (Figure 3E). PC markers (MEF2C, PEGFRβ, RGS5) were highly expressed in the early stages but down-regulated in the later stages, while SMC markers (ACTA2, MYL9, TAGLN) showed opposite tendency (Figure 3E).

**Figure 3.**
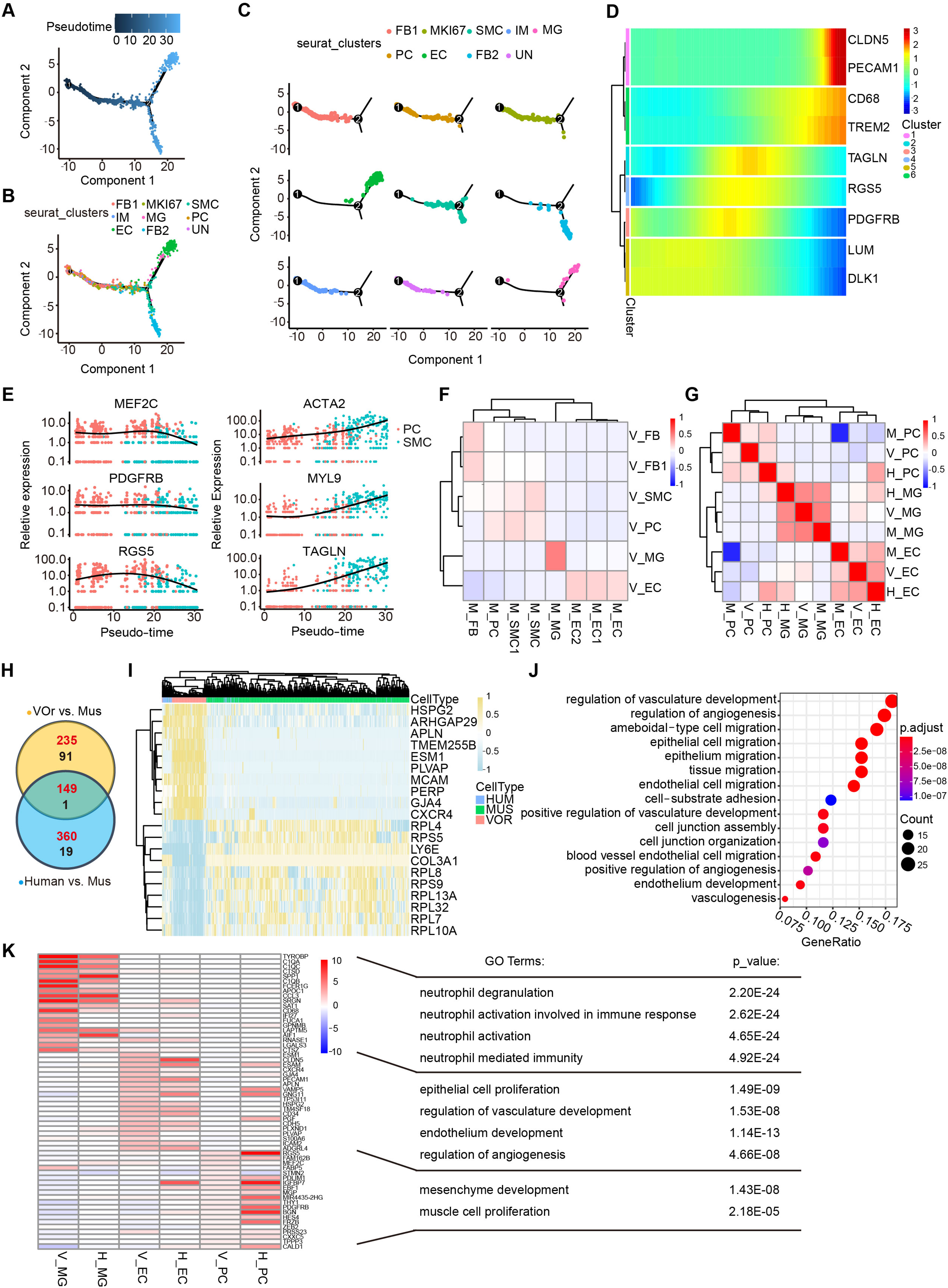
Cell fate trajectory analysis in VOrs and the comparison with cell types *in vivo*. (**A**) Single-cell trajectories by monocle analysis showing developmental stage of the VOrs. (**B**) Clusters in UMAP showing trajectory track. (**C**) Developmental trajectory of indicated cell clusters in VOrs. (**D**) Heatmap showing the expression level of the main cell type-specific markers with pseudo-time. (**E**) Expression of markers in SMC and PC with pseudo-time. (**F**) Correlation analysis of cell clusters (EC, MG, PC, SMC, FB) between VOrs and mouse brain. V, data from VOrs; M, data from mouse. (**G**) Correlation analysis of cell clusters (EC, MG, PC) among VOrs, mouse and human brain single-cell data. V, data from VOrs; M, data from mouse; H, data from human. (**H**) Venn diagram showing the DEGs in EC clusters for VOr and human samples compared to mouse samples. Red for up-regulated genes, black for down-regulated genes. (**I**) Heat-map showing the top enriched DEGs in the EC cluster for VOrs samples compared to mouse sample (fold change > 1.25 and p < 0.05). (**J**) GO analysis of the 149 up-regulated DEGs in (h) (p value < 0.1 and FDR < 0.05). (**K**) Top 20 marker genes for VOrs in the main clusters (EC, PC, MG) (fold change > 1.25 and p < 0.05) compared to human sample, with significantly pathways by GO analysis (p value < 0.1 and FDR < 0.05). V, data from VOrs; H, data from human.

In order to determine to what extent the VOrs resembled the brain vessels *in vivo*, we analyzed two accessible datasets for comparison. First, we compared the VOrs and mouse cerebrovascular scRNA-seq data (He et al., 2018; Vanlandewijck et al., 2018), and found that the molecular features of the five major cell types, including FB, SMC, PC, EC, and MG, in VOrs were similar to the counterparts of mouse cerebrovascular system (Figure 3F). We then referred to a dataset of scRNA-seq from eight adult and four embryonic human cortexes, which clustered a small number of vascular cell types, including EC, PC and MG (Polioudakis et al., 2019). We analyzed the correlation of these three clusters in VOrs, mouse and human samples together, and found that VOrs and human showed stronger correlation in EC and PC clusters, while MG showed the highest consistency across all three datasets (Figure 3G). This result further confirmed the presence of brain specific MG cells in VOrs culture system with the introduction of neurotrophic factors. Thus, VOr is an appropriate model for the analysis of human cerebrovascular development *in vitro*.

Next we analyzed differentially expressed genes (DEGs) in ECs between VOrs and mouse, human and mouse, respectively, and found that a big fraction of genes up-regulated were overlapped between the two sets of comparisons (Figure 3H). Remarkably, most of the top DEGs between VOrs and mouse groups showed similar tendency in human samples (Figure 3I), suggesting the high similarity of ECs in VOrs compared to that in human samples *in vivo*. The gene ontology (GO) analysis showed that the shared DEGs between VOr vs. mouse and human vs. mouse pairs were related to the angiogenesis pathway, suggesting that human vascular development may be more complex and robust than that of mouse (Figure 3J). We also analyzed DEGs within PCs, and found that, the majority of top changed genes in VOrs compared with mouse samples were also present in DEGs of human vs. mouse pair comparison (Figure 3-figure supplement 1D and E). To further validate that VOrs can faithfully mimic the process of vascular development *in vivo*, the expression of marker genes of three major cell types from VOrs (EC, PC, MG) were compared with that of human samples. As shown in Figure 3K, marker genes of VOrs were also highly expressed in the same cell types of human samples, further indicating the similarity of corresponding cell type. Recently, Lu et al demonstrated that some *in vitro* induced brain vessel cells lacked functional attributes of ECs but were more related to the neuroectodermal epithelial lineage-induced brain microvascular endothelial cells (Epi-iBMEC) (Lu et al., 2021). We performed principal componant analysis (PCA) for ECs in VOrs and other 28 datasets from this study, including the primary ECs, induced ECs (iEC), and Epi-iBMECs, and found that ECs in VOrs showed a clear disparity from Epi-iBMECs but higher similarity to EC lineage (Figure 3-figure supplement 1F). The top and bottom loading genes showed separate endothelial and epithelial cell type identities in these datasets (Figure 3-figure supplement 1G), and VOrs exhibited strong endothelial cell properties (Figure 3G). These results indicate that the VOr model can be used to recapitulate cerebrovascular development *in vitro*.

### Generation of fusion vascularized brain organoids

Having established the VOrs, we decided to generate vascularized brain organoids by using co-culture strategy. For this purpose, we established the induction system of brain organoids from human H9 ESCs according to the methods reported previously (Lancaster and Knoblich, 2014; Mariani et al., 2012; Ou et al., 2020), with some modifications (Figure 4-figure supplement 1A). Cerebral organoids at different developmental stages were stained with neural progenitor markers PAX6 and p-VIM, the proliferation marker KI67, intermediate progenitor marker TBR2, young neuron marker DCX (doublecortin), mature neuron marker TUJ1, and the cortical layer markers (TBR1, CTIP2, SATB2, REELN), and the results indicated that the brain organoids were well induced (Figure 4-figure supplement 1B-E). As expected, CD31^+^ ECs were not observed in this induction system (Figure 4-figure supplement 1F). After the step of neural ectoderm induction, EBs with neuroepithelial (NE) property were co-embedded with VPs in one Matrigel droplet, and then cultured under the condition of VOrs maturation with the medium containing N2 and B27 (Figure 4A). For better invasion of vessels into the developing BOrs, we put two VP bodies in both sides of one NE body (Figure 4A). After co-culture for different days, VOrs labelled by GFP gradually wrapped BOrs and finally formed a fused vasculature and brain organoids (fVBOrs) by D40 (Figure 4B). Whole-mount staining of the fVBOrs showed that DCX-labelled neurons were enwrapped by invaded vessels labeled by CD31 (Figure 4C).

**Figure 4.**
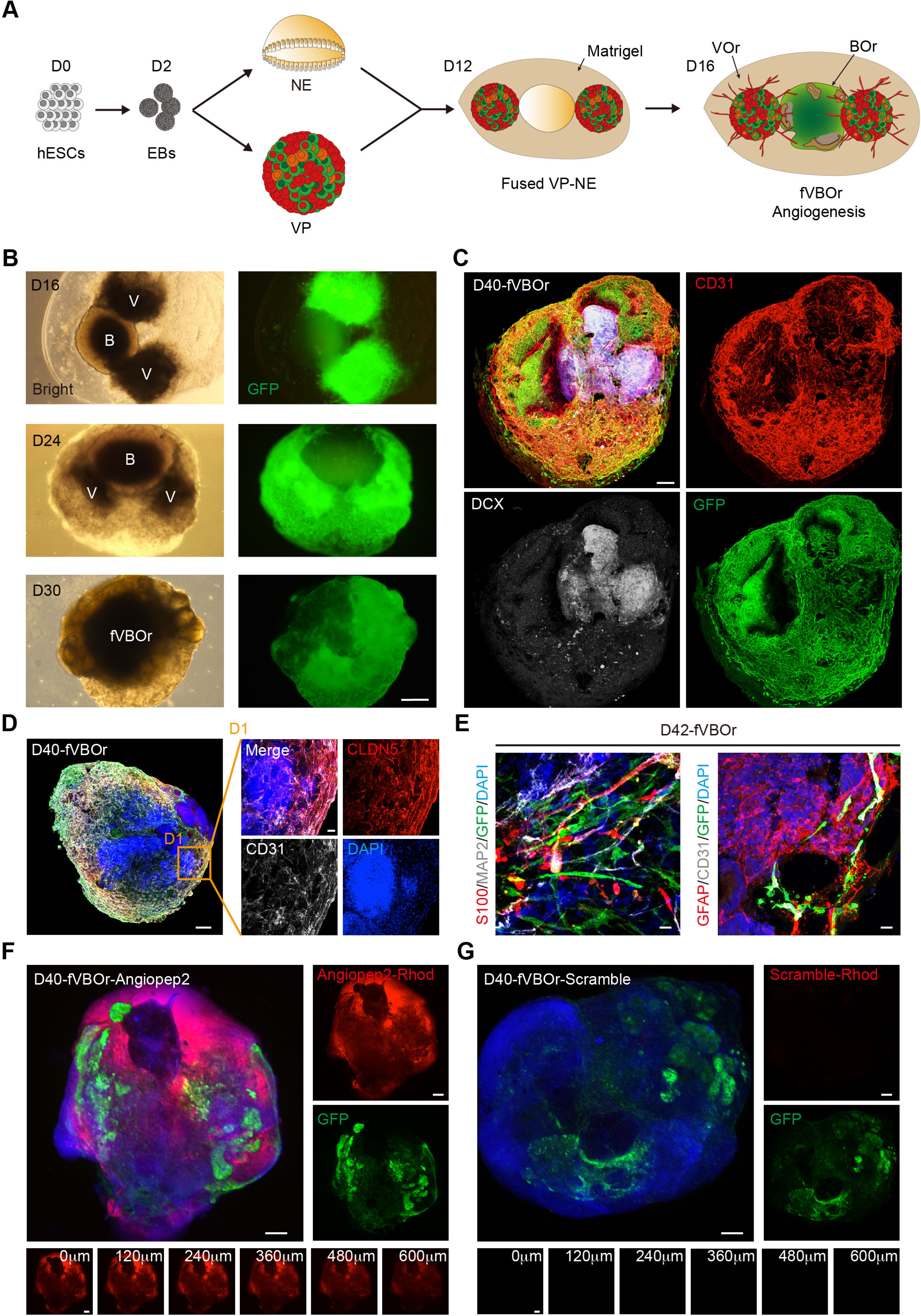
Generation of fVBOrs with BBB structure. (**A**) Schematic view of the method for generating fVBOrs. EBs, embryonic bodies; NE, neuroepithelium; VP, vascular progenitor; VO, vessel organoid; BOr, brain organoid; fVBOr, fusion vascularized brain organoid. (**B**) fVBOrs at different developmental stages. Scale bar, 500 μm. V, VOr; B, BOr. (**C**) Immunostaining of CD31 and DCX for labeling vessels and neurons, respectively, in D40 fVBOrs. Scale bar, 200 μm. (**D**) Immunostaining of CLDN5 for labeling tight junctions in fVBOrs. Scale bar, 200 μm. D1, enlarged area. (**E**) Immunostaining for markers of astrocytes (S100/GFAP), neurons (MAP2), endothelial cells (CD31) and vessel structures (GFP) in fVBOrs. Orange arrows indicate astrocytes end feet. Scale bar, 20 μm. (**F, G**) Confocal fluorescence images showing the transport of rhodamine-labeled angiopep-2 (Angiopep-2–Rhod), rhodamine–scramble peptide (Scramble–Rhod) in fVBOrs. Scale bar, 200 μm. Bottom, z-stack images of rhodamine signals.

Supported by pericytes and astrocytes, the brain microvascular ECs form a particularly tight layer called the blood-brain-barrier (BBB), which selectively controls the flow of substances into and out of the brain by forming complex intercellular tight junctions and protects the brain from harmful substances (Augustin and Koh, 2017; Chow and Gu, 2015; Lippmann et al., 2012; Sweeney et al., 2019). To determine whether the fVBOrs developed BBB-like features, we examined the expression of the tight junctions proteins Claudin5 (CLDN5) and ZO-1 (Figure 4D and Figure 4-figure supplement 2A and B), and the efflux transporter p-Glycoprotein which helps the recycling of small lipophilic molecules diffused into ECs back to the blood stream (Augustin and Koh, 2017; Lippmann et al., 2012) (Figure 4-figure supplement 2C). Notably, stronger CLDN5 signals were observed in BOr regions in contact with vessels, suggesting the appearance of tight junctions-like structures (Figure 4-figure supplement 2A). Furthermore, astrocyte-like cells labelled by S100 or GFAP were also observed in fusion organoids, forming neurovascular unit-like structures composed of CD31/GFP-labelled vascular structures and MAP2-labelled neurons (Figure 4E). The presence of neurovascular unit structure in fusion organoids was also confirmed by Transmission Electronic Microscopy (TEM), which showed the endothelial cell basement membrane enclosed by pericytes and tightly contacted by end-feet of astrocytes (Figure 4-figure supplement 2E).

Next, we examined the functionality of BBB in the fVBOrs, by measuring the permeability of molecules with different BBB penetration capability (Bergmann et al., 2018; Cho et al., 2017; Dai et al., 2018; Xu et al., 2019). The selectivity of BBB was determined by incubating fVBOrs with rhodamine-labelled Angiopep-2, a peptide capable of permeating through BBB selectively (Bergmann et al., 2018; Cho et al., 2017). We found that Angiopep-2 exhibited strong signals in the fVBOrs, but scrambled peptides displayed no detectable signal (Figure 4F and G). The z-stack images showed that the intensity of Angiopep-2 signals decreased from the surface to the inner of fVBOrs (Figure 4F, bottom). In contrast to fusion organoids, the BOrs alone showed much weaker Angiopep-2 signals (Figure 4-figure supplement 2E and F). Taken together, these results indicate that fVBOrs have developed BBB structures with selective permeability.

### Microglia cells in fVBOrs are responsive to immune stimuli

It is generally believed that MGs are developed from the yolk-sac progenitors, which then populate in the developing brain to regulate neurogenesis and neural circuit refinement (Kaur et al., 2017; Mosser et al., 2017; Salter and Stevens, 2017). Indeed, in the unguided cerebral organoids, spontaneous MG can emerge (Ormel et al., 2018), probably due to the presence of residue mesodermal progenitors (Quadrato et al., 2017). However, the functional investigation is limited due to the variable and inconsistent batch effects. Based on the scRNA-seq and staining results suggesting the presence of MG in VOrs, we decided to explore the possibility of introducing these MG-like cells into brain organoids using fusion strategy. To this end, we first determined the molecular features of MG in VOrs. As shown in Figure 5A, the MG identity was confirmed by the expression of specific marker genes *AIF1* (the gene encoding IBA1) and *CD68*. The GO analysis showed that the functions of MG markers were mainly concentrated on the pathways of immune and inflammatory responses (Figure 5B). During the VOr culture, the expression of MG marker genes, such as *AIF1* or *TMEM119*, gradually increased (Figure 5C), indicating again the MG induction. In line with this notion, D40 VOrs exhibited increased abundance of MG with amoeboid-like morphology, compared to D25 MG mostly with round morphology (Figure 5D and E).

**Figure 5.**
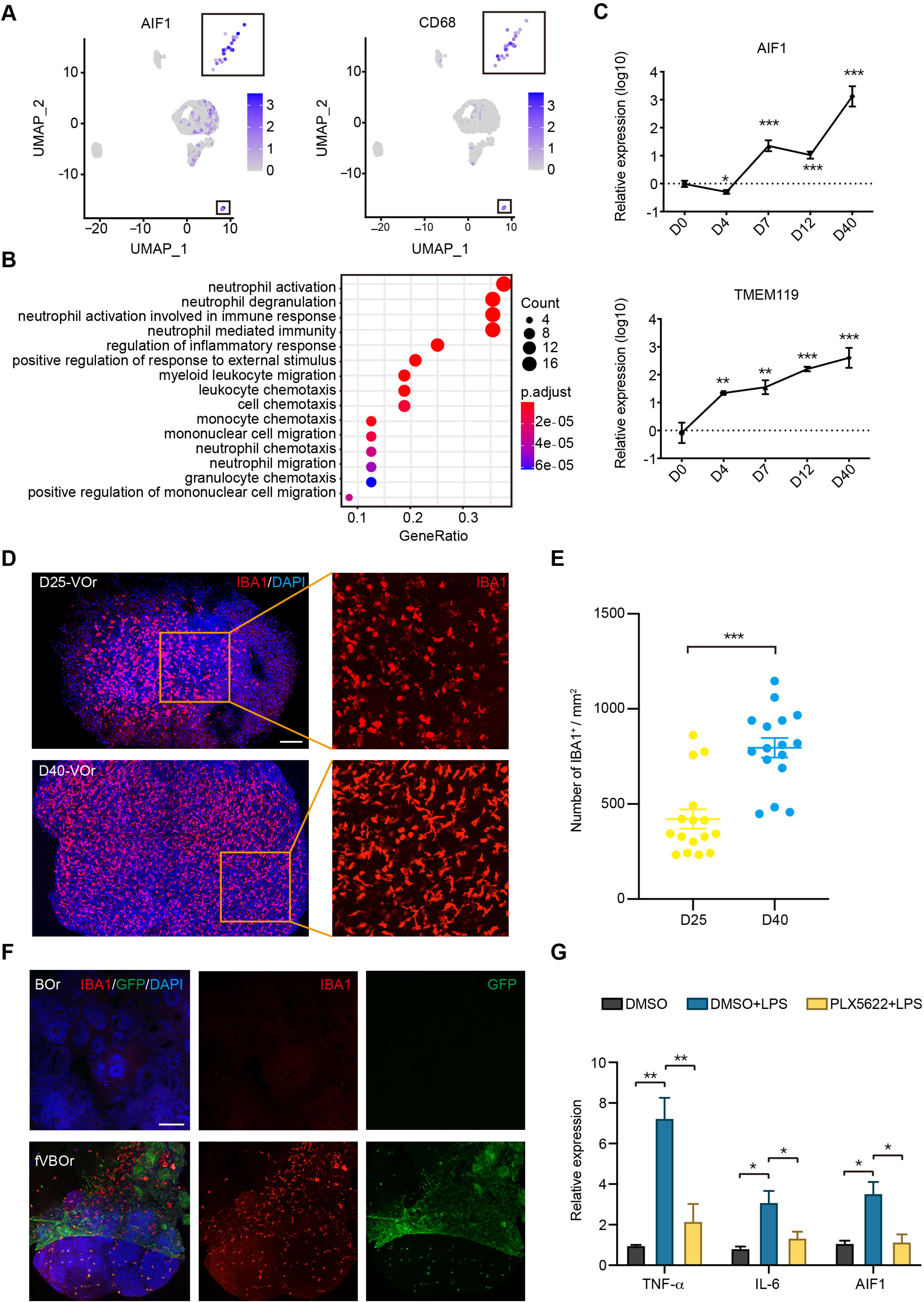
Microglial cells in fVBOrs. (**A**) UMAP plot showing single-cell expression pattern of microglial specific markers in VOr. Relative expression level is plotted from gray to blue colors. (**B**) GO analysis of microglial cell marker genes (p value < 0.1 and FDR < 0.05). (**C**) qPCR analysis for expression of microglial markers AIF1 and TMEM119 in developing VOrs. Data are presented as mean ± SEM (n = 3 independent experiments with 6–7 organoids in each group at indicated time point). **p< 0.01, ***p< 0.001. (**D**) Immunostaining of IBA1 for labeling microglial cells in D25 and D40 VOrs. Scale bar, 200 μm. (**E**) Quantification of the IBA1^+^ cell number in D25 and D40 VOrs. n = 16. Students t-test, ***p < 0.001. (**F**) Immunostaining of IBA1 for labeling microglial cells in BOrs and fVBOrs, respectively. Scale bar, 200 μm. (**G**) qPCR analysis for the expression of indicated genes in D40 fVBOrs treated with LPS (500 ng/ml, MCE, HY-D1056) without or with PLX5622 2 μM (MCE, HY-11415) using DMSO as vehicle control. Relative expression was normalized to GAPDH. n = 3 independent experiments with 8-10 organoids in each group. One-way ANOVA, *p < 0.05, **p < 0.01.

We next examined whether MG could migrate from VOrs into BOrs after fusion. We found large amount of IBA1^+^GFP^+^ MG-like cells in the neural part of fVBOrs, whereas BOrs alone had no MG-like signal (Figure 5F). Thus, fVBOrs also contained MG-like cells, besides vasculatures. Then, we determined the responsiveness of these MG-like cells to the treatments used to diminish or activate microglia to verify the cell identity. First, we treated D30 VOrs with PLX5622, a selective inhibitor of CSF1R (colony stimulating factor 1 receptor), which was used to ablate MG in mice (Huang et al., 2018). After treatment for 7 days with 2 μM PLX5622, the IBA1-labelled cells were almost completely gone, and the ablation effect lasted for at least 3 days in the absence of PLX5622 (Figure 5-figure supplement 1A and B). Likewise, the PLX5622 treatment also depleted MG-like cells in D40 fVBOrs (Figure 5-figure supplement 1C). These results further confirmed the identity of MG-like cells in VOrs and fVBOrs. Next, fVBOrs were treated with 0.5 μg/ml lipopolysaccharide (LPS) for 72 h to induce inflammatory response. The LPS stimulation caused marked increase in the expression of inflammatory factors TNFα and IL-6 (Figure 5G). Interestingly, the expression levels of TNFα or IL-6 were attenuated in PLX5622-treated fusion organoids (Figure 5G), suggesting the involvement of MG-like cells in LPS-induced immune response. These results support the conclusion that MG-like cells possess responsive ability to immune stimuli.

### Increased neural progenitors in the fusion organoids

It has been shown that vasculature acts as a critical niche that helps maintain the survival and stemness of neural progenitors, and EC-derived soluble factors might contribute to this function (Delgado et al., 2014; Ottone et al., 2014; Shen et al., 2004). Prompted by these information, we compared neurogenesis patterns in fVBOr and BOr. Interestingly, the fVBOrs exhibited marked increase in the thickness of neuroepithelial rosettes compared to BOrs at the same corresponding stages (D25) (Figure 6A and B). In line with this notion, the density of neural progenitors (NP) marked by PAX6 or mitotic NPs marked by phospho-vimentin (P-VIM) also increased in fusion organoids (Figure 6C-E). However, the density of DCX-labelled differentiated neurons or TBR1-labelled early born cortical neurons had no difference at the observation period (Figure 6F and G; Figure 6-figure supplement 1A and B). These results suggest that the factors produced by VOrs might promote the proliferation of NPs after fusion with BOrs, with little effect on neuronal differentiation. In the classical brain organoid culture system, the inner cells are extremely vulnerable to limited accessibility to the trophic factors in culture medium. In line with this notion, BOrs at D40 showed abundant apoptotic cells expressing cleaved caspase 3 (c-CASP3) in the central regions, whereas the apoptotic cells were markedly reduced in fVBOrs (Figure 6-figure supplement 1C-E). Thus, the fusion organoids generated in this work can be used to study interactions among multiple cell types during brain development.

**Figure 6.**
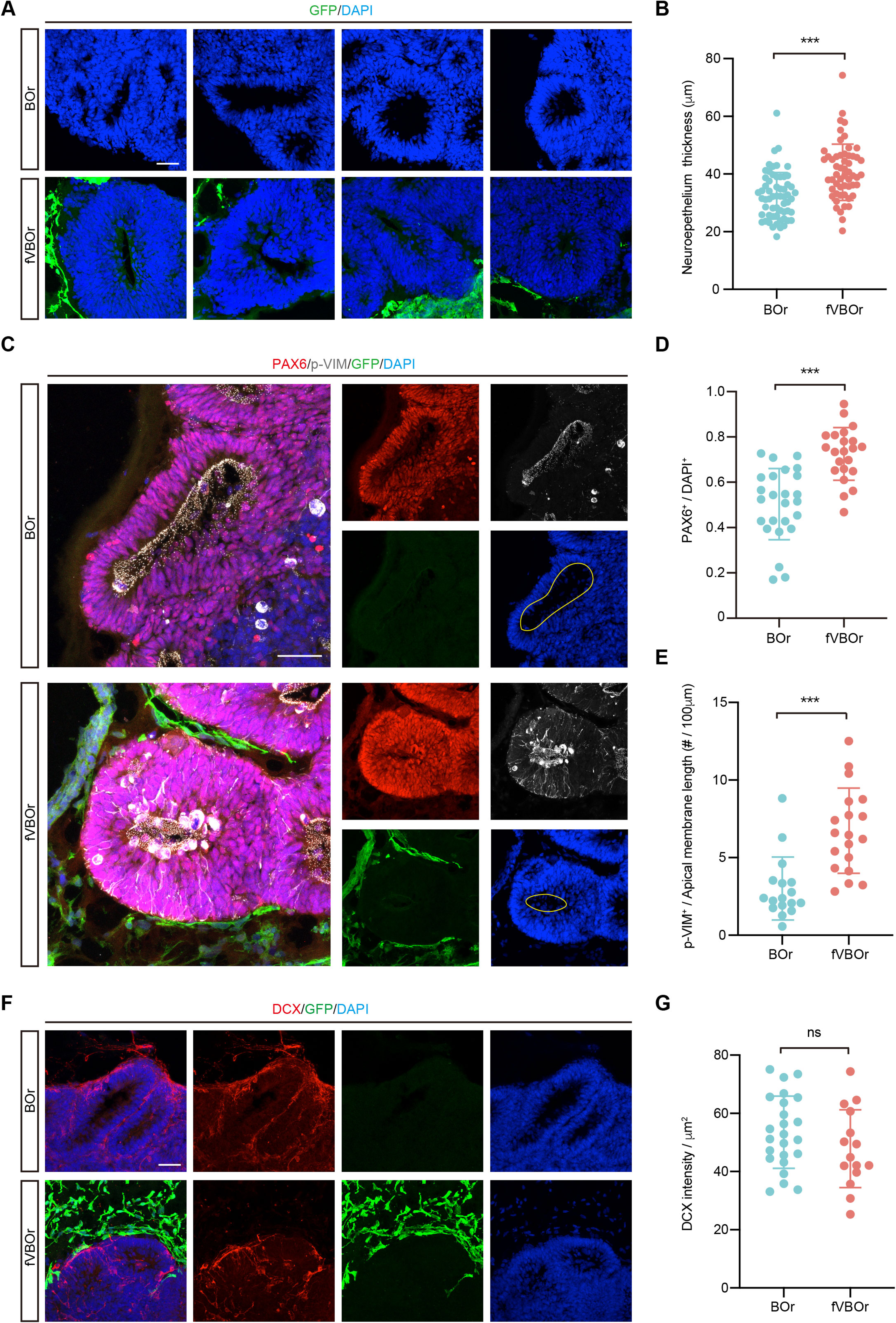
Increased neurogenesis in fVBOrs. (**A**) Immunostaining for DAPI showing the neuroepithelium rosettes of BOrs and fVBOrs at D25. Scale bar, 50 μm. (**B**) Quantification of neuroepithelium thickness of BOrs and fVBOrs. Data are presented as mean ± SEM (BOrs: n = 60 rosettes from 7 organoids; fVBOrs: n = 55 rosettes from 6 organoids). Two-tailed Student’s *t*-test. ***p < 0.001. (**C**) Immunostaining for PAX6 and p-VIM in VZ-like area of BOrs and fVBOrs at D25. Scale bar, 50 μm. Apical membrane was showed in yellow circle. (**D, E**) Quantification of the density of PAX6^+^ (**D**) and the density of p-VIM^+^ cells per 100 μM apical membrane length (**E**) in BOrs and fVBOrs. Data are presented as mean ± SEM (PAX6: n = 25 rosettes from 4 organoids; p-VIM: n = 23 rosettes from 4 organoids). Two-tailed Student’s *t*-test. ***p < 0.001. (**F**) Immunostaining for DCX in BOrs and fVBOrs at D25. Scale bar, 50 μm. (**G**) Quantification of the intensity of DCX in BOrs and fVBOrs. Data are presented as mean ± SEM (BOrs: n = 24 rosettes from 3 organoids, fVBOrs: n = 15 rosettes from 4 organoid). ns, no significant difference (p = 0.308, two-tailed Student’s *t*-test).

## Discussion

The prevalent approaches for brain organoid induction start from neuroectoderm induction using BMP and/or Wnt signaling inhibitors, or neural induction medium to treat embryoid bodies, followed by brain regional differentiation, and neural maturation (Giandomenico and Lancaster, 2017; Kadoshima et al., 2013; Lancaster and Knoblich, 2014; Lancaster et al., 2013; Qian et al., 2016). Although this system is powerful, it lacks vascularization rendering it impossible to recapitulate vascular-neural interactions and model related diseases. Here, we developed an approach for generating neural-specific vessel organoids by initial transient mesoderm induction, sequential VP and EC induction, followed by treatments with neurotrophic reagents. Then, we established an integrated vascularized brain organoid model by fusing the brain and vessel organoids. This vascularization strategy considered the compatibility to mesodermal and ectodermal linages and led to generation of brain organoids with complex tubular vessels, functional neurovasculature units, as well as microglia responding to immune stimulation.

The fVBOr model generated in this study provides a possibility to analyze the process of brain angiogenesis and complex interactions between vasculatures and neural cells. It has been shown previously that brain vascularization is regulated by neural progenitors (Matsuoka et al., 2017), and, on the other hand, vasculatures promote neurogenesis and oligodendrocyte precursor migration (Tata et al., 2016; Tsai et al., 2016). Notably, we found that in fVBOrs, only vasculatures close to or located in the BOrs expressed tight junctions markers, including CLDN5 and ZO-1 (Figure 4-figure supplement 2A and B), consistent with the idea that BBB maturation is regulated by neural cues (Lippmann et al., 2012). The effects of vasculatures on neurogenesis were also found in fVBOrs, which exhibited increased pool of neural progenitors and reduced apoptosis (Figure 6C-E; Figure 6-figure supplement 1C-E). The reduction in apoptotic cells was also seen in grafted BOrs with the invasion of host blood vessels (Mansour et al., 2018; Shi et al., 2020). It is conceivable that the vascularized brain organoids developed in this study may provide a feasible platform for the study of human brain development, vasculature-related diseases or pharmaceutical interventions, which need to pass the BBB barrier.

Several studies have tried to generate BBB-like structures *in vitro*, by culturing PSC-derived ECs (Lippmann et al., 2012; Qian et al., 2017) or co-culturing PSC-derived cells without or with primary cells in 2D system (Appelt-Menzel et al., 2017; Canfield et al., 2017) or 3D system (Bergmann et al., 2018; Cho et al., 2017). Although some features of the BBB were reproduced in these studies, the tube-like structures of blood vessels were lacking. The vessel organoid model we have established showed intense vascular network with characteristics and specificity of human cerebral vessels based on transcriptomic analysis and functional assay. It has been demonstrated that the combination of PSC-derived tissue-specific progenitors or relevant tissue samples with ECs and mesenchymal stem cells can generate vascularized organs (Takebe et al., 2015; Takebe et al., 2013), but different derivation of vascular ECs and the brain from distinct germ layers limits the vascularization in BOrs initialized from neuroectoderm induction. Indeed, we failed to observe any vascular structures in the brain organoids generated using the prevalent approach (Figure 4-figure supplement 1F). Although the fVBOr model has shown branched vessels, it still lacks active blood flow. One possible approach to solve the problem is to combine microfluidic techniques and organoid cultures and make an “organ-on-a-chip”, which may authentically mimic the vascular environment.

The blood vessels in the brain are not only required for oxygen and nutrient supply, but also involved in regulating neurogenesis. The fusion organoids allow for recapitulating early developmental processes like vasculogenesis and angiogenesis, as well as the integration of microglia. Unlike BBB-like structures generated in other studies that combined various mature cell types (Cho et al., 2017; Lippmann et al., 2012; Qian et al., 2017), the ones generated in our study were directly induced from pluripotent stem cells, which resembled the developmental processes *in vivo*. Although the factors that defined the identity of microglia cells in VOrs are not clear, their migration into the brain organoids resembled the extra-embryonic originality. It is known that astrocytes are essential components of the neurovascular units (Abbott et al., 2006), their involvement in the maturation of BBB-like units or immune surveillance awaits for further investigation.

## Materials and Methods

### hESCs culture

H9 human embryonic stem (H9-hES) cell line was purchased from iMedCell. H9-hES-EGFP was generated by introducing the CAG-EGFP DNA fragment into the genome locus ROSAβgeo26 (ROSA26) using the CRISPR/Cas9 method. Both H9-hES and H9-hES-EGFP cells were cultured and passaged as previously described (Ou et al., 2020). Cells were cultured on hESC-Matrigel (Corning) coated dishes in mTeSR1 (STEMCELL) medium with the addition of bFGF (4 μg/ml, STEMCELL). The culture medium was half-replaced every day and then cells were passaged every 5 days using passage reagent ReLeSR (STEMCELL).

### Generation of human brain organoid

*h*ESC clones were dissociated into single cells with Accutase (STEMCELL), then cells were re-suspended in mTeSR1 medium containing 10 μM Y27632 (STEMCELL), and seeded into the lipidure-coated (NOF CORPRATION) V-bottom 96-well plate (Thermo) with 7000 cells per aggregate, 150 μl per well to form EBs. On day 2, the culture medium was replaced by the ectodermal induction medium (DMEM/F12 (Life/Invitrogen) containing 20% (v/v) Knockout Serum Replacer (Gibco), 1% (v/v) MEM-NEAA (Gibco), 3.5 μl/l β-mercaptoethanol (Sigma-Aldrich), 1% (v/v) Glutamax (Gibco), 2.5 μM dorsomorphine (Tocris) and 2 μM A83-01 (Tocris)). On day 4, the ectodermal induction medium was half-replaced. On day 6, the EB medium was replaced by the neural induction medium (DMEM/F12 containing 1% (v/v) N2 supplement (Life/Invitrogen), 1% (v/v) MEM-NEAA, 1% (v/v) Glutamax, 1 μg/ml heparin (Sigma-Aldrich,), 10 μM SB431542 (Selleck) and 200 nM LDN193189 2HCL (Selleck)) and lasted for 6 days. The neural induction medium was half-renewed every other day. On day 12, the EBs were embedded into growth factor-reduced Matrigel droplet (Corning), as described previously (Lancaster and Knoblich, 2014). And the culture medium was replaced by the differentiation medium (50% (v/v) DMEM/F12 and 50% (v/v) Neurobasal medium (Life/Invitrogen) containing 0.5% (v/v) N2 supplement, 0.5% (v/v) B27 supplement without vitamin A (Life/Invitrogen), 3.5 μl/l β-mercaptoethanol, 250 μl/l Insulin (Sigma-Aldrich), 1% (v/v) Glutamax and 0.5% (v/v) MEM-NEAA, 1% (v/v) Antibiotic-Antimycotic (Gibco)). After 4 days, the differentiation medium was replaced by the maturation medium (50% (v/v) DMEM/F12 and 50% (v/v) Neurobasal medium containing 0.5% (v/v) N2 supplement, 0.5% (v/v) B27 supplement (Life/Invitrogen), 3.5 μl/l β-mercaptoethanol, 250 μl/l Insulin, 1% (v/v) Glutamax and 0.5% (v/v) MEM-NEAA, 1% (v/v) Antibiotic-Antimycotic). Then, the organoids were transferred into a shaker in the 5% CO_2_ incubator at 37°C for maturation, and the medium was renewed every 3-4 days.

### Generation of human vessel organoid

H9-hES-GFP clones were dissociated into single cells with Accutase, then the cells were re-suspended in the mTeSR1 medium containing 10 μM Y27632 and seeded into the lipidure-coated V-bottom 96-well plate with 9000 cells per aggregate, 150 μl per well to form EBs. On day 2, the culture medium was replaced by the mesodermal induction medium (APEL2 (STEMCELL) with 6 μM CHIR99021 (Selleck)). On day 4, the mesodermal medium was replaced by endothelial induction medium (APEL2 with 50 μg/ml VEGF (STEMCELL), 25 μg/ml BMP4 (R&D, 314-BP) and 10 μg/ml bFGF). On day 7, medium was changed into MV2 medium (Promocell) with 50 μg/ml VEGF for the maturation of endothelial cells, and the medium was renewed every other day. From day 12, the EBs were embedded into Matrigel droplets and cultured with VEGF-containing (20 μg/ml) neural differentiation medium, as that used for brain organoid culture.

### Fusion of vascular brain organoids

To generate the fusion organoids, two VOr EBs and one BOr EB were collected and then embedded together into one Matrigel droplet (25 μl) on day 12. The two VOr EBs were put on two sides of the BOr EB, and pipette tips could be used to adjust the shape and site of the three EBs. The following steps were the same as the non-fusion BOr EBs with the addition of 20 μg/ml VEGF.

### Immunofluorescence

The collected organoid samples were fixed in 4% paraformaldehyde (PFA) at 4°C overnight, and then washed 3 times with PBS, dehydrated in 30% sucrose at 4°C for 24-48 hr. Then, organoids were embedded in O.C.T (Sakura) and cryosectioned into 30 μm-thick slices. The sectioned slices were boiled in citrate-based antigen retrieval buffer for 10 min, followed by cooling for over 60 min. Slices were washed in PBS for 3 times and incubated in 0.3% TritonX-100 (Sigma-Aldrich,) at room temperature (RT) for 30 min, blocked with 5% BSA (Sigma-Aldrich) in 0.1% TritonX-100 at RT for 1 hr, incubated with the primary antibody at 4°C for over 48 hr, followed by washes with PBS and incubation with the secondary antibody at 4°C overnight. Secondary antibodies were: AlexaFluor 488, 555, 594, or 647-conjugated donkey anti-mouse, -rabbit, -rat or -chicken IgG (Invitrogen, all used at 1:1000 dilution). DAPI (beyotime, 1:2000 dilution) was used to mark cell nuclei. Stained sections were mounted with mounting medium and stored at 4°C before imaging. All images were acquired by confocal imaging systems.

For whole-mount staining, the organoid samples were fixed in 4% PFA at 4°C overnight, and then washed for 3 times with PBS, followed by the incubation in 0.5% TritonX-100 at RT for 1 hr. After blocking with 5% BSA in 0.1% TritonX-100 at RT for 1 hr, organoids were incubated with primary antibodies at 4°C for over 48 hr, washed with PBS, and then incubated with secondary antibodies at 4°C for over 48 hr. The stained organoids were washed by PBS for 3 times before confocal imaging.

### Quantitative PCR (qPCR)

The total RNA of 3-4 organoids was extracted using the RNeasy Plus Micro Kit (Qiagen), followed by reverse transcription to generate cDNA with GoScript Reverse Transcription Kit (Promega). Quantitative PCR was performed by using the Agilent Mx3000P qPCR system with the 2×SYBR Green qPCR Master Mix (Bimake). Relative mRNA expression was determined by the delta cycle time with human GAPDH as the internal control in data normalization. Primer sequences were as follows:

TNF-α: forward, 5’- CACAGTGAAGTGCTGGCAAC -3’, reverse, 5’-AGGAAGGCCTAAGGTCCACT -3’;
IL-6: forward, 5’- TTCCAAAGATGTAGCCGCCC-3’, reverse, 5’-ACCAGGCAAGTCTCCTCATT-3’;
GAPDH: forward, 5’-TCGGAGTCAACGGATTTGGT-3’, reverse, 5’-TTCCCGTTCTCAGCCTTGAC-3’;
IBA1: forward, 5’-AAACCAGGGATTTACAGGGAGG-3’, reverse, 5’-GGGCAGATCCTCATCACTGC-3’;
TMEM119: forward, 5’-GAGGAGGGACGGGAGGAG-3’, reverse: 5’-GACCAGTTCCTTGGCGTACA-3’;
NANOG: forward, 5’-CAATGGTGTGACGCAGAAGG-3’, reverse: 5’-TGCACCAGGTCTGAGTGTTC-3’;
OCT4: forward, 5’-CTCGAGAAGGATGTGGTCCG-3’, reverse, 5’-TGACGGAGACAGGGGGAAAG-3’;
PECAM1: forward, 5’-AGACGTGCAGTACACGGAAG-3’, reverse, 5’-TTTCCACGGCATCAGGGAC-3’;
VE-Cadherin: forward, 5’-CGCAATAGACAAGGACATAACAC-3’, reverse, 5’-GGTCAAACTGCCCATACTTG-3’;
VWF: forward, 5’-CCCGAAAGGCCAGGTGTA-3’, reverse, 5’-AGCAAGCTTCCGGGGACT-3’;
VEGFR2: forward, 5’-GAGGGGAACTGAAGACAGGC-3’, reverse, 5’-GGCCAAGAGGCTTACCTAGC-3’;
VEGFR1: forward, 5’-AACGTGGTTAACCTGCTGGG-3’, reverse, 5’-AGTGCTGCATCCTTGTTGAGA-3’;
PDGFR: forward, 5’-ATCAGCAGCAAGGCGAGC-3’, reverse, 5’-CAGGTCAGAACGAAGGTGCT-3’.

### Flow cytometry

VOrs were dissociated with Trypsin solution as described in single-cell dissociation. After re-suspension in staining buffer, about 1×10^6^ single cells for each group were incubated with Alexa 647-labeled CD31 antibody (1:1000 dilution, BD) for 30 min. The results were analyzed by using FlowJo software. Single cells isolated from BOrs were used as the negative control.

### LDL-uptake assay

hESC (D0) and VOrs (D4-D40) were washed with PBS for 3 times and then incubated with 10 μg/ml Ac-LDL (Yeasen) in MV2 medium for 4 hr at 37°C. The samples were washed for 3 times with PBS, before confocal imaging using 20×objective lens.

### BBB penetrating assay

fVBOrs were collected in 35-mm dishes and washed with PBS for 3 times, followed by incubation with 5 μM angiopep (TAMRA-TFFYGGSRGKRNNFKTEEY) or control scrambled (TAMRA-GNYTSRFEREYGKFNKFGT) peptides in neuronal maturation medium for 3 hr at 37°C. Then, the organoids were washed with PBS, fixed in 4% PFA containing DAPI to stain the cell nuclei, and then imaged using 1.25× objective lens.

### Transmission Electron Microscope (TEM) analysis

Cultured fVBOrs were washed with DPBS (Life/Invitrogen) and cut into 1 mm × 1 mm small pieces. Firstly samples were fixed with 4% paraformaldehyde overnight and then were pre-fixed with 2.5% glutaraldehyde (SPI, USA), in PBS for 12 hr. After washing with PBS, samples were post-fixed with 1% OsO4 (TED PELLA) for 2 hr at 4°C. Next, followed by dehydrated in an ascending gradual series (30%–100% (v/v)) of ethanol, and embedded in epoxy resin (Pon812 kit, SPI, USA). The embedded samples were initially cut into about 500 nm-thick sections, inspected by stained with toluidine blue (Sinopharm), and finally sectioned into 70-nm by Leica EM UC7. Then sections were double-stained with uranyl acetate (SPI, USA) and lead citrate (SPI, USA), followed by observation with a Transmission Electron Microscopy (Talos L120C) at an acceleration voltage of 80 kV.

### Single cell dissociation and 10x genomics chromium library construction

Organoids were dissociated using the methods as described previously (Thomsen et al., 2016). Briefly, 8-10 organoids were collected and washed by DPBS (Life/Invitrogen,) and cut into small pieces, followed by the incubation with 2 mL trypsin solution (Ca^2+^/Mg^2+^-free HBSS (Life/Invitrogen) with 10 mM HEPES (Sigma-Aldrich), 2 mM MgCl_2_, 10 μg/ml DNase Ι (Roche), 0.25 mg/ml trypsin (Sigma-Aldrich), PH 7.6) for 30 min at 37°C. Then the samples were quenched with 4 ml Quenching Buffer (440 ml Leibovitz L-15 medium (Thermo) with 50 ml ddH_2_O, 5 ml 1 M HEPES (pH 7.3-7.4), 10 μg/ml DNase I, 100 nM TTX (TOCRIS), 20 μM DNQX (TOCRIS), and 50 μM DL-AP5 (TOCRIS), 5 ml 100× Anti-Anti, 2 mg/ml BSA, 100 μg/ml trypsin inhibitor (Sigma-Aldrich)), subjected to centrifugation in 220×g for 4 min at 4°C, and re-suspended with 2 ml Staining Medium (440 mL Leibovitz L-15 medium with 50 ml ddH_2_O, 5 ml 1 M HEPES (pH 7.3-7.4), 1 g BSA, 100 nM TTX, 20 μM DNQX, and 50 μM DL-AP5, 5 ml 100× Anti-Anti, 20 ml 77.7 mM EDTA (pH 8.0)), filtered through a 40 micron cell filter (Falcon), centrifuged again in 220×g for 4 min at 4°C, then the cells were re-suspended in 5 ml DPBS with 1% BSA. Dissociated cells were re-suspended at a concentration of 500 cells/μl. cDNA libraries were generated following the guidelines provided by 10x Genomics, Inc. Briefly, dissociated cells were partitioned into nanoliter-scale Gel Bead-In-Emulsions (GEMs), and then subjected to reverse transcription, cDNA amplification and library construction, with individual cell and transcript barcoded.

### Single-cell RNA-Seq data analysis

Cellranger software was used for mapping raw data to the human genome (version hg38 (v1.2.0)). And then data were processed with Seurat (v3.0) under R (v3.5.2) environment. For quality control, cells expressing less than 200 genes or more than 7000 genes were removed, and genes expressed in less than 3 cells were excluded for the following analysis. VST (variance-stabilizing transformation) was used for searching for the highly-variable genes and the top 2,000 genes were chosen to do the downstream analysis. The cells were clustered and reduced into the UMAP space by principal component analysis (PCA), with 1st to 15th principal components (PCs). Differentially expressed genes (DEGs) in each cluster were identified by more than 1.25 fold change and p value < 0.05 with Wilcoxon Rank Sum test. GO enrichment analysis was carried out by ‘ClusterProfiler’ and org.Hs.eg.db in R software program with p value < 0.1 and FDR (false discovery rate) < 0.05 considered as statistical significance.

Monocle (v2.10.1) package was used to analyze the developmental trajectory of cell types. The monocle object was constructed from the Seurat object. After normalization and variance estimation, we calculated the mean and dispersion values and chose the genes whose mean expression were >0.1. Then cells were dimensional reduced and clustered. DEGs with >1.25 fold change and p value < 0.05 (two-side t test) were used for downstream analysis. The cell trajectory plots were produced by running ‘reduceDimension’ and ‘orderCell’ function with defined option.

Human neocortical scRNA-seq data (Polioudakis et al., 2019) (phs001836) and mouse brain vascular scRNA-seq data (He et al., 2018; Vanlandewijck et al., 2018) (GSE98816) were downloaded for the correlation analysis. The pearson correlation coefficient calculated by the intersect genes between different data sets was used for correlation analysis. DEGs were set with 1.25 fold change and p value <0.05 by limma (v3.38.3) package.

## Data availability

Single cell RNA sequencing transcriptome data supporting this study have been deposited in NCBI Sequence Read Archive (SRA) repository (https://www.ncbi.nlm.nih.gov/sra) with accession number SRP338043 (VOR: SRR15992286; VOR2:SRR15992285). All software used is open and freely available.

## Acknowledgments

This study was partially supported by the National Key Research and Development Program of China (2021ZD0202500), National Natural Science Foundation of China (32130035 and 92168107 to Z.G.L, 31871034 to X.C.J), the Frontier Key Project of the Chinese Academy of Sciences (QYZDJ-SSW-SMC025), Shanghai Municipal Science and Technology Projects (2018SHZDZX05, 201409001700), and National Key R&D Program of China (2017YFA0700500). We are grateful to the Multi-Omics Core Facility, Molecular Imaging Core Facility and Molecular and Cell Biology Core Facility at the School of Life Science and Technology, ShanghaiTech University for providing technical support.

## Author Contributions

X.Y.S., X.C.J., designed and conducted the experiments; Y.J.C contributed unpublished reagents; X.Y.S., X.C.J., Y.L., P.M.Z., performed the experiments; X.Y.S., P.M.Z., L.B.S., performed the single-cell analysis; Y.L., J.W. participated in sample preparation; X.Y.S and Z.G.L wrote the paper. Z.G.L supervised the whole work.

---

**Figure 1—figure supplement 1.**
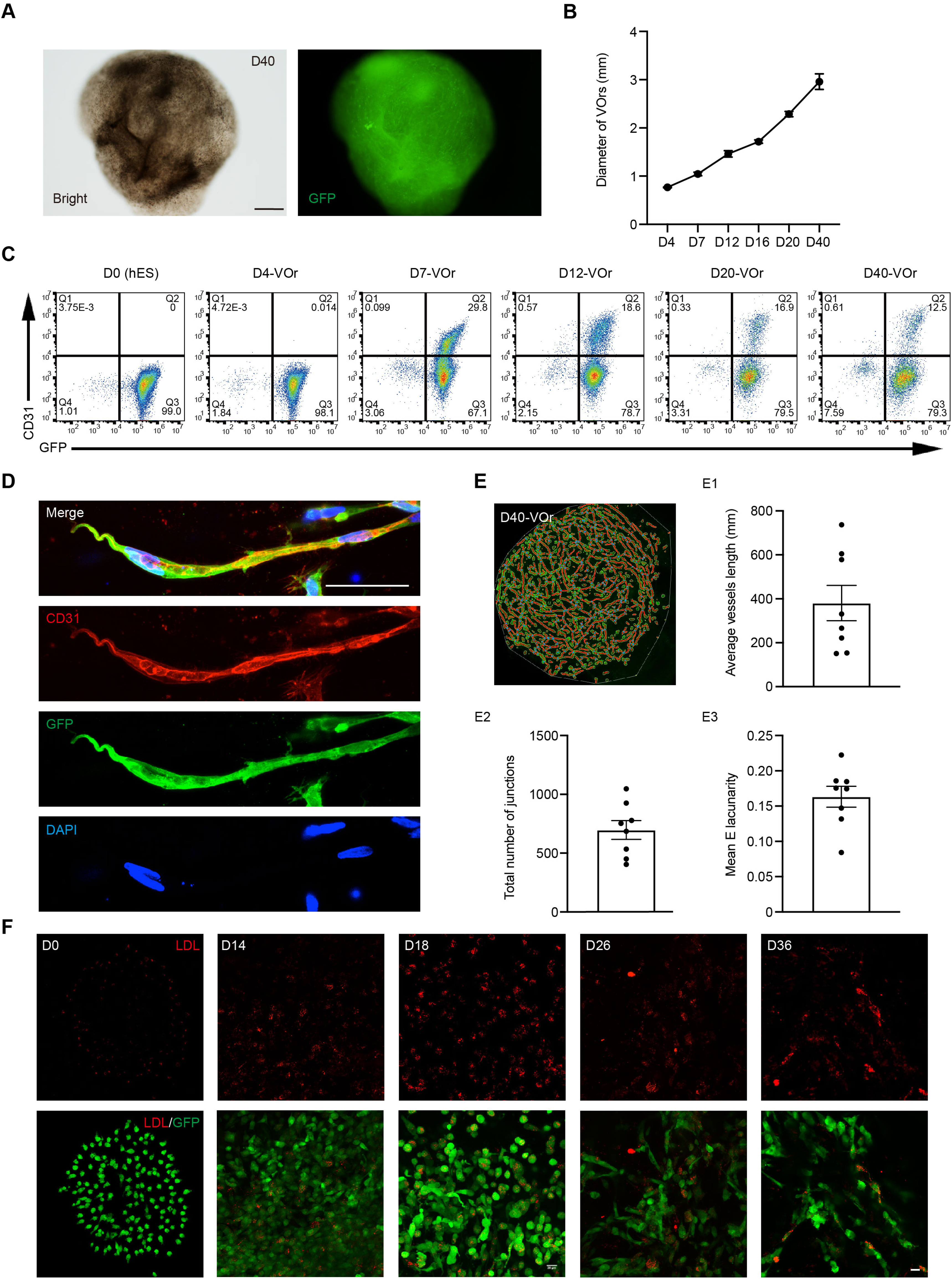
VOrs recapitulate human vessel development. (**A**) Morphological appearance of VOrs at D40. Scale bar, 500 μm. (**B**) Quantification of the VOr diameter from D4 to D40. Data are mean ± SEM of 10-21 organoids in each time points. (**C**) Flow cytometry plots of temporal development of CD31^+^ GFP^+^ cells in differentiating VOrs. (**D**) Immunostaining of GFP and CD31 for the vascular angiogenesis structures. Scale bar, 20 μm. (**E**) Quantification of the average vessel length (**E1**), total number of junctions (**E2**) and mean E lacunarity (**E3**) for D40 VOrs. (**F**) Up-take of acetylated low-density lipoprotein (DiI-Ac-LDL) of VOrs at indicated time points. Scale bar, 20 μm.

**Figure 2—figure supplement 1.**
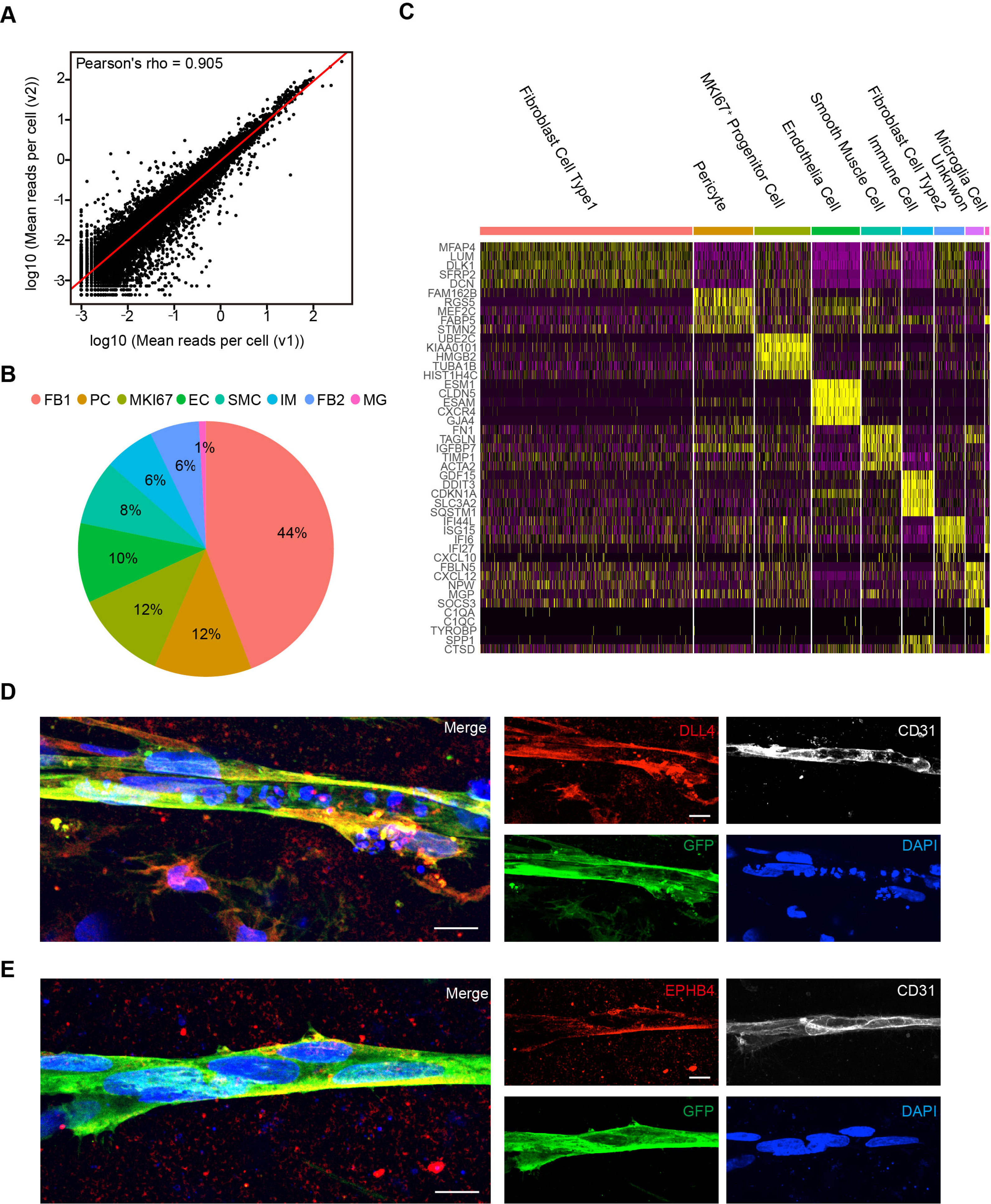
Cell type analysis for VOrs by scRNA-seq and immunostaining. (**A**) Correlation analysis of scRNA-seq data from two batches of VOr samples. (**B**) Proportions of cell types among all the cells from VOrs. (**C**) Heatmap showing the top five most enriched genes for each cell type. (**D**) Immunostaining of DLL4 for labeling the arterial endothelial cells in VOrs. Scale bar, 10 μm. (**E**) Immunostaining of EPHB4 for labeling the venous endothelial cells in VOrs. Scale bar, 10 μm.

**Figure 3—figure supplement 1.**
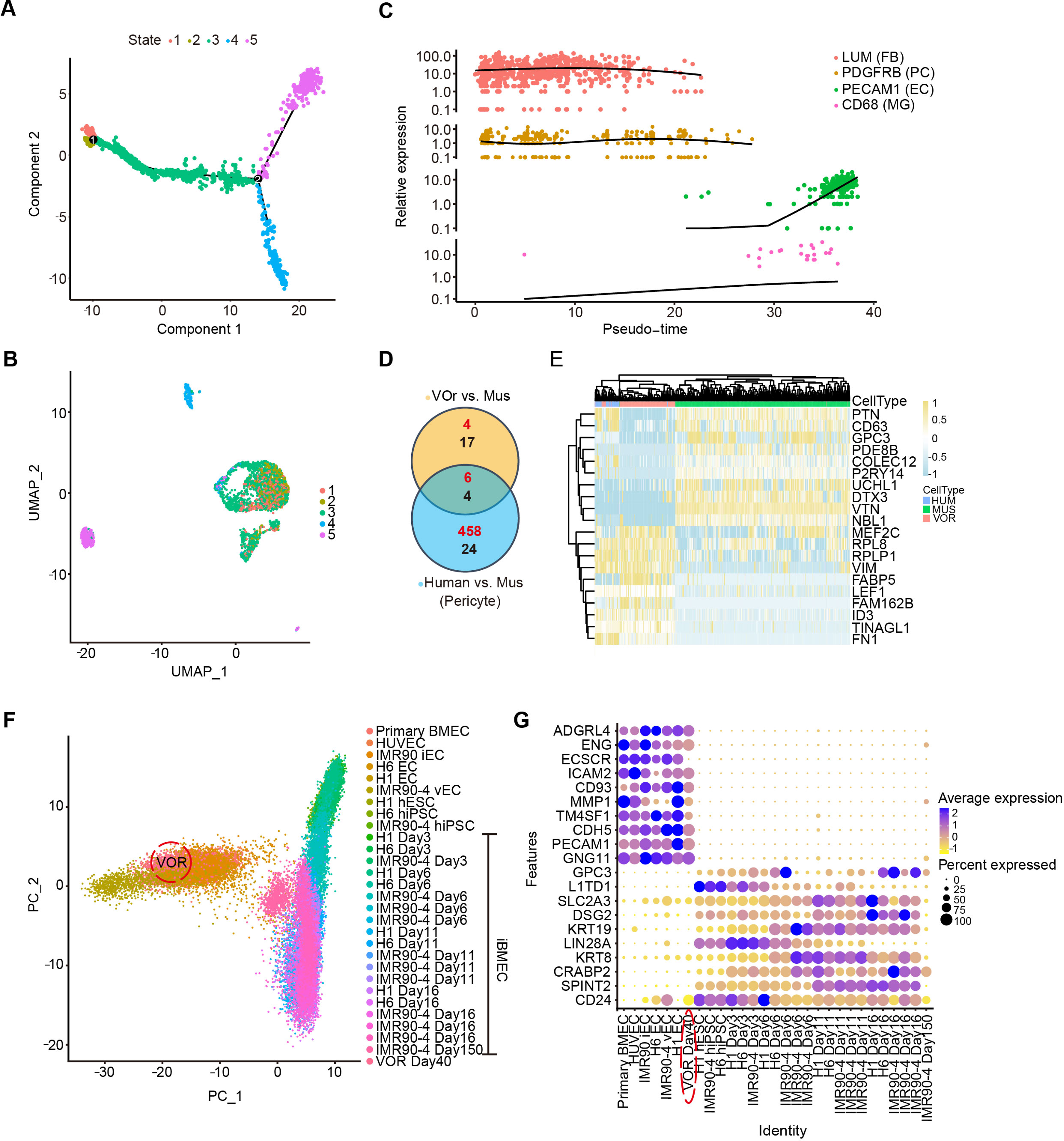
Cell types in VOrs are similar to that of human samples *in vivo*. (**A**) Trajectory analysis showing five main developmental stages. (**B**) UMAP plot showing five developmental stages of VOrs. (**C**) Expression of markers in four main clusters with pseudo-time. (**D**) Venn diagram showing the DEGs in PC clusters for VOr and human samples compared with mice samples. Red for up-regulated genes, black for down-regulated genes. (**E**) Heat-map showing the top enriched common DEGs in the PC cluster for VOrs samples compared to mouse sample (fold change > 1.25 and p < 0.05). (**F**) Principal component analysis for relative relationship of 29 distinct cell samples from published datasets, including primary ECs, iECs (induced ECs), and Epi-iBMECs (neuroectodermal epithelial lineage-induced brain microvascular endothelial cells) at various days of differentiation and hPSCs (human pluripotent stem cells). ECs in VOrs are highlighted to emphases the similarity to ECs rather than iBMECs. (**G**) Heat-map illustrating differences in expression levels of both endothelial- and epithelial-specific genes in all cell samples from (**F**). The VOR sample (circled) shows more EC-specific gene expression rather than epithelial-specific expression.

**Figure 4—figure supplement 1.**
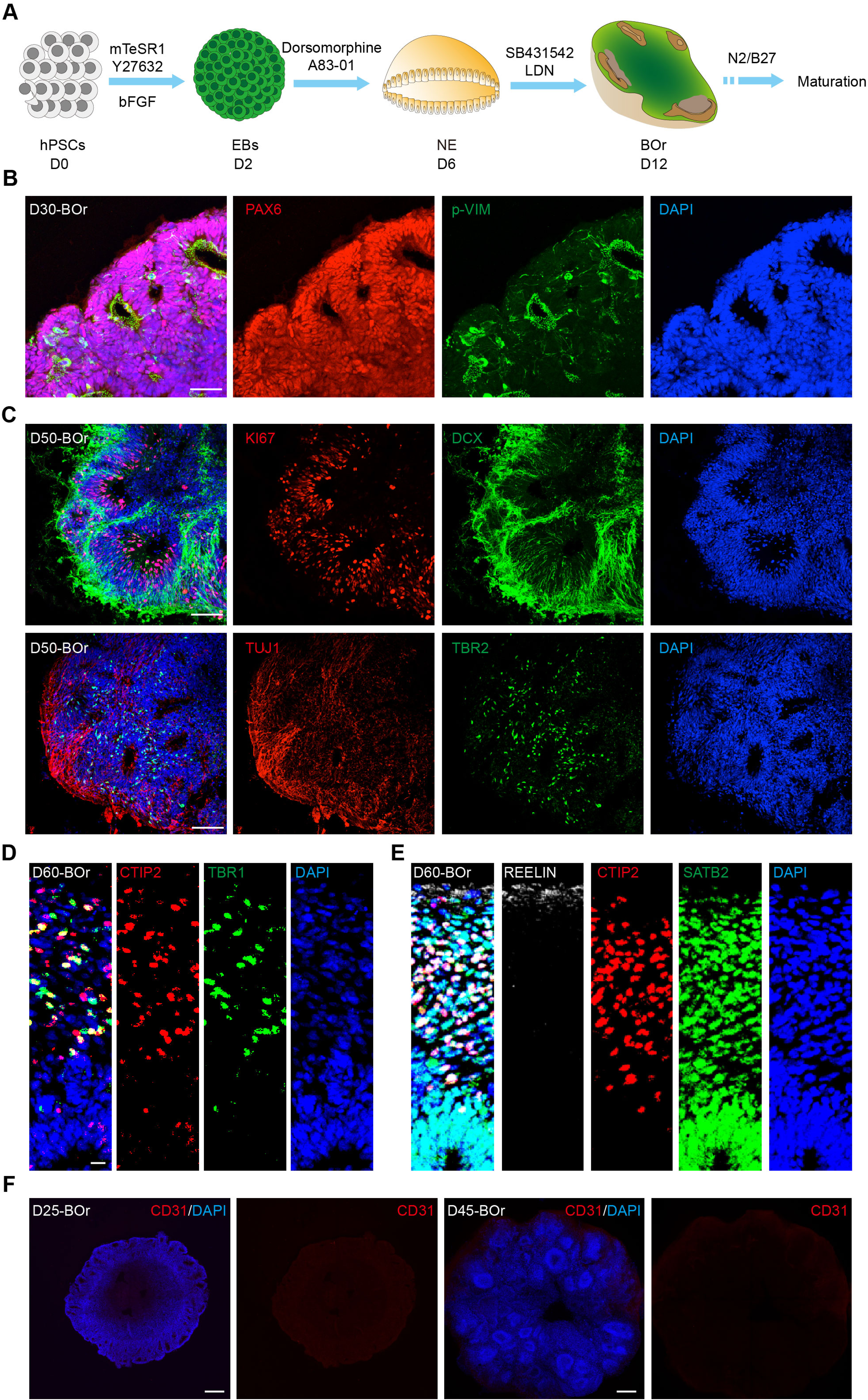
Generation of human brain organoids. (**A**) Schematic workflow for generating BOrs from hESC. EBs, embryonic bodies; NE, neuroepithelium; BOr, brain organoid. (**B**) Immunostaining for cortical progenitor markers PAX6 and p-VIM in BOrs at D30. Scale bar, 50 μm. (**C**) Immunostaining for young neuron marker DCX, proliferation marker KI67, intermediate progenitor marker TBR2 and neuron marker TUJ1 in D50 BOrs. Scale bar, 50 μm. (**D, E**) Immunostaining for cortical layer markers TBR1, CTIP2, SATB2 and REELIN in D60 BOrs. Scale bar, 50 μm. (**F**) Immunostaining for EC marker CD31 in BOrs at D25 and D45. Scale bar, 200 μm.

**Figure 4—figure supplement 2.**
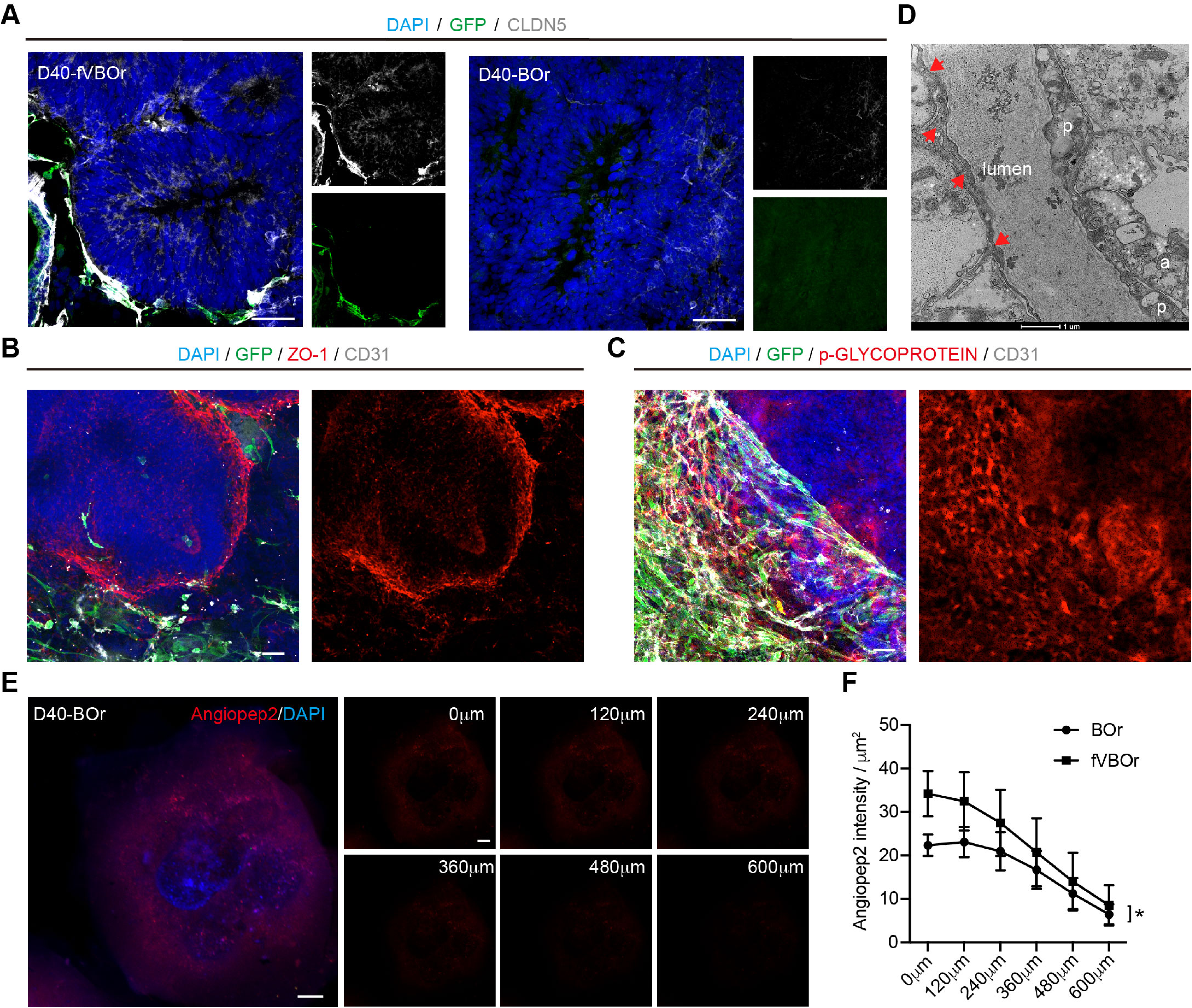
BBB-like structures in fVBOrs. (**A**) Fluorescence image showing the expression of CLDN5 in fVBOrs and BOrs respectively. Scale bar, 50 μm. (**B**) Fluorescence image showing the expression of tight junctions marker ZO-1 in fVBOrs at D40. Scale bar, 50 μm. (**C**) Fluorescence image showing the expression of the efflux transporter p-Glycoprotein in fVBOrs at D40. Scale bar, 50 μm. (**D**) TEM of vascular and BBB structure in D80-fVBOr. Arrows: basement membrane; p: pericyte; a: end-foot of astrocyte. Scale bar, 1 μm. (**E**) Confocal z-stack images of rhodamine-labeled angiopep-2 (Angiopep-2–Rhod) in D40 BOrs. (**F**) Quantification for the intensity of Angiopep-2–Rhod across different z-stack images of fVBOrs compared with BOrs. Data are presented as mean ± SEM (n = 3 organoids in each sample). *p < 0.05, paired t-test.

**Figure 5—figure supplement 1.**
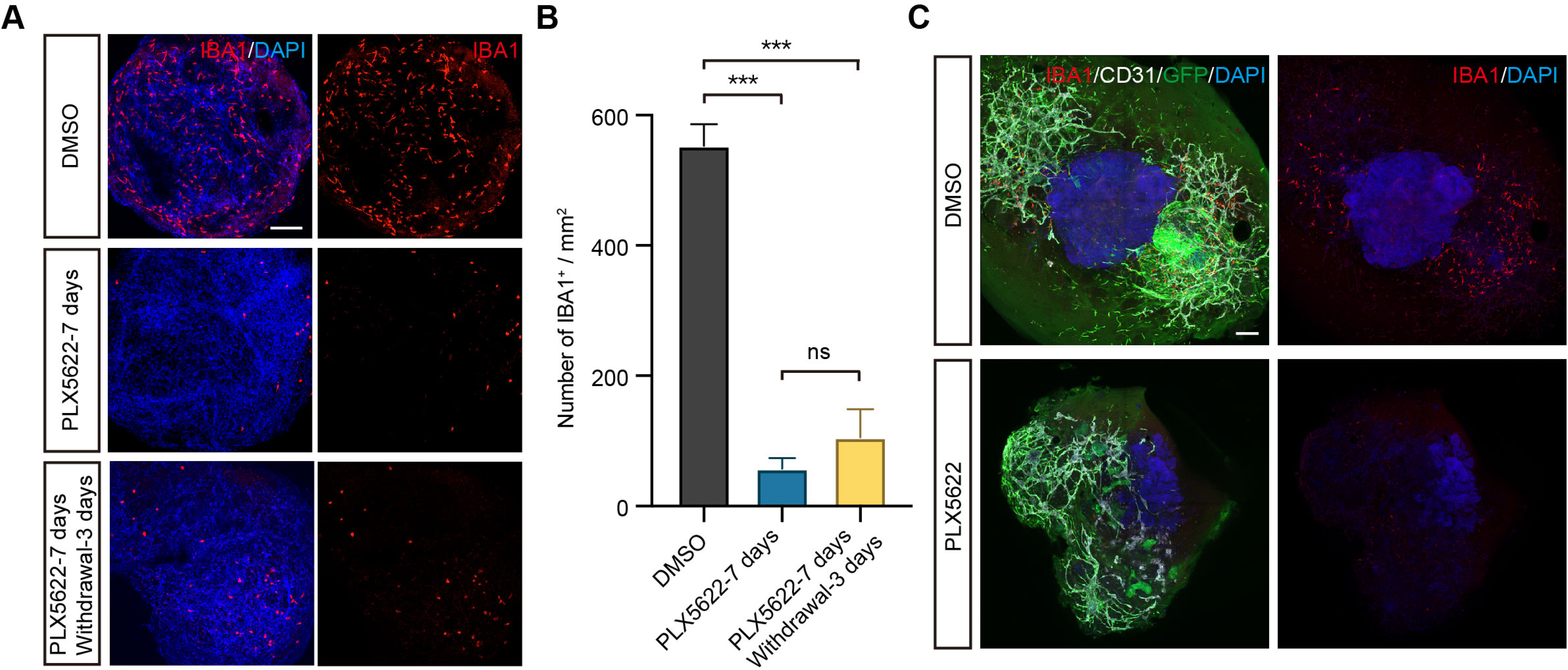
PLX5622 ablates MGs in VOrs. (**A**) IBA1 staining in VOrs showing ablation of microglial cells after treatment with 2 μM PLX5622 for 7 days, and drug withdraw for 3 days. Scale bar, 200 μm. (**B**) Quantification of microglial cell numbers in VOrs with indicated treatments. Data are shown as mean ± SEM (n = 3 independent experiments with 8-10 organoids in each group) One-way ANOVA, ***p < 0.001, ns, no significant difference. (**C**) Ablation of microglia in fVBOrs treated with 2 μM PLX5622 for 7 days. Scale bar, 200 μm.

**Figure 6—figure supplement 1.**
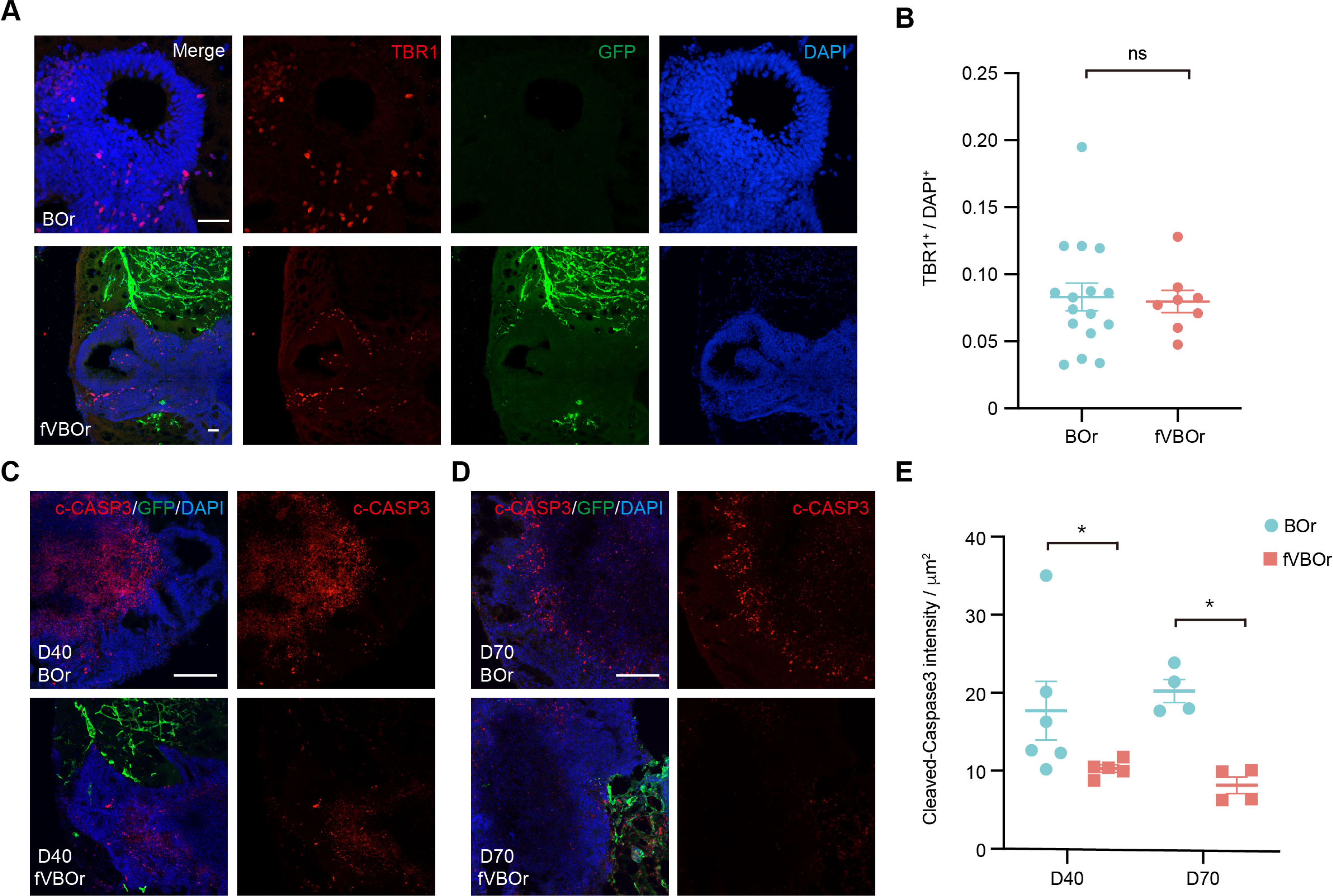
Reduced apoptotic areas in fVBOrs. (**A**) Immunostaining for early born deep-layer neuronal marker TBR1 in BOrs and fVBOrs at D40. Scale bar, 50 μm. (**B**) Quantification of TBR1^+^ cells in BOrs and fVBOrs (BOr: n = 16 rosettes from 6 organoids, fVBOr: n = 8 rosettes from 3 organoids). Data are presented as mean ± SEM (two-tailed Student’s *t*-test, p = 0.8340). (**C, D**) Staining of cleaved-caspase3 (c-CASP3) in BOrs or fVBOrs at D40 (**C**) and D70 (**D**). Scale bar, 200 μm. (**E**) Quantification of the cleaved-caspase3 intensity in BOrs and fVBOrs at D40 and D70, respectively. Data are presented as mean ± SEM of chosen fields of 5 organoids in each sample. *p< 0.05 (p = 0.0303 for D40, p = 0.0286 for D70, two-tailed Student’s *t*-test).

**Movie Supplement 1**

PBS fluid was microinjected into the vessel-like lumen in D40 VOr with continuous pressure, showing liquid flow and vessel wall expansion without leakage. Electrode inner diameter: 500 nm.

**Supplementary File 1.** Key resources table

